# Harnessing the Power of Large Language Models (LLMs) to Unravel the Influence of Genes and Medication on Biological Processes of Wound Healing^*^

**DOI:** 10.1101/2024.03.26.586862

**Authors:** Jayati H. Jui, Milos Hauskrecht

## Abstract

Recent advancements in Large Language Models (LLMs) have ushered in a new era for knowledge extraction in the domains of biological and clinical natural language processing (NLP). In this research, we present a novel approach to understanding the regulatory effects of genes and medications on biological processes central to wound healing. Utilizing the capabilities of Generative Pre-trained Transformer (GPT) models by OpenAI, specifically GPT-3.5 and GPT-4, we developed a comprehensive pipeline for the identification and grounding of biological processes and the extraction of such regulatory relations. The performances of both GPTs were rigorously evaluated against a manually annotated corpus of 104 PubMed titles, focusing on their ability to accurately identify and ground biological process concepts and extract relevant regulatory relationships from the text. Our findings demonstrate that GPT-4, in particular, exhibits superior performance in all the tasks, showcasing its potential to facilitate significant advancements in biomedical research without requiring model fine-tuning.

## 1 Introduction

Wound healing is a multifaceted biological process that plays a crucial role in restoring the integrity and functionality of injured tissues, involving a series of sequential and overlapping phases like inflammation, proliferation, and remodeling ^1,2^. The cellular and biological events of wound healing are regulated by the synergistic influence of numerous genes, proteins, and potential drug targets. Knowledge about the regulatory effects of genes and medications on the underlying mechanisms of wound healing is dispersed across a vast array of scientific publications, which presents a challenge to achieve a comprehensive and consolidated understanding of the topic. However, with the rise of conditional and large language models (LLM), there’s been a notable shift within the NLP realm towards harnessing the capabilities of LLMs to solve specialized downstream NLP tasks. In this study, our aim is to utilize the advanced capabilities of the Generative Pretrained Transformer (GPT) models developed by OpenAI, specifically GPT-3.5 and GPT-4, to uncover the regulatory effects of genes and medications on the biological processes involved in wound healing by parsing scientific texts.

Identifying biological functions in text through Named Entity Recognition (NER) poses significant challenges. The description of biological functions often varies due to the complex linguistic structures and semantic nuances inherent in natural language, thus complicating the task of recognizing these phrases as unified concepts for conventional NER systems. Additionally, the occurrence of intervening words within phrases may cause NER systems to erroneously interpret them as separate entities. For instance, the biological function “fibroblast proliferation” might be expressed in several ways, such as “proliferating fibroblasts”, “proliferation of fibroblasts”, or “fibroblast cell proliferation” etc. In a previous work, we suggested using dependency rule-based methods for entity phrase augmentation and merging to enhance the recognition of biological functions ^3^. Nonetheless, relying exclusively on a set of predefined rules has its limitations, particularly in capturing the broad and dynamic range of natural language expressions.

Linking biological function concepts identified in natural text to established knowledge bases like UMLS or Gene Ontology (GO)^†^ enhances standardization and minimizes ambiguity in scientific terminology. Grounding these concepts not only validates the accuracy of the information extracted but also supports computational biology tasks by adhering to globally recognized concepts. However, traditional biological knowledge graphs, such as Hetionet, DRKG, and Bioteque, often miss detailed regulatory relationships between gene, medications and biological processes, limiting their effectiveness for drug repurposing ^4–6^. By identifying and incorporating these regulatory relations into existing knowledge graphs, their utility for drug repurposing can be significantly improved.

In this study, we introduce a comprehensive relation extraction framework that utilizes the advanced analytical abilities of GPT-3.5 and GPT-4. However, the primary challenge with these LLMs is the potential inability to incorporate the latest updates from the continuously evolving Gene Ontology (GO) database. By integrating the Retrieval Augmented Generation (RAG) technique with the GPT models, we can enhance their capabilities to identify and ground biological process concepts accurately. RAG improves the annotation by pulling in the latest information from external GO databases in real time, ensuring responses of these LLMs are based on the most current knowledge. This method also reduces the well-known risk of hallucination. Furthermore, we utilize both GPT-3.5 and GPT-4 to discern the regulatory impacts of genes and medications on biological functions.

In this study, we used a curated corpus of fully annotated titles derived from scientific article titles that depict such relationships. We evaluate the ability of GPT3.5 and GPT-4 to identify and standardize concepts of biological processes, as well as its performance in extraction of regulatory relations. Our key contributions in this study are summarized as follows:

- We utilize GPT-3.5 and GPT-4 to identify concepts of biological processes in natural language texts and standardize these concepts to the Gene Ontology (GO) database.
- We employ both GPT models for the extraction of regulatory effects of genes and medications on biological processes.
- We conduct a comprehensive assessment of the effectiveness of the GPPT models across the three tasks, comparing their performance with a baseline established by a traditional NLP tool for knowledge extraction.

## 2 Background

With the emergence of LLMs, many of the latest works in relation extraction shifted towards prompt-based structured prediction. Most of the LLM-based studies primarily concentrate on prompt-based few-shot in-context learning and utilizing synthetic data produced by LLMs to enhance the accuracy of relation extraction. In a recent work, Wadhwa et. al. assessed modern LLMs, specifically GPT-3 and Flan T5 (Large), for relation extraction and found that GPT-3 is comparable to SOTA models with minimal examples ^7^. Using a distillation method with Chain of Thought (CoT) explanations from GPT-3 to fine-tune Flan-T5, they achieved leading performance, suggesting LLMs as a potential standard baseline for RE. In a similar work, Xu et. al. delved into the potential of in-context learning and data generation using GPT-3.5 for few-shot relation extraction, with enhancements from task-related instructions and schema-constrained data generation ^8^. Xu and colleagues proposed a two-stage self-training framework using synthetic data from LLMs for relation extraction, addressing data scarcity and domain differences. Their method showed significant performance improvement, setting new benchmarks for low-resource relation extraction ^9^. Yuan and colleagues explored ChatGPT’s proficiency in zero-shot temporal relation extraction using three prompt techniques, finding its performance lagging behind supervised methods and highly dependent on prompt design ^10^. Their work highlighted the limitations of ChatGPT in terms of consistency and long-dependency temporal inference. Some recent studies have harnessed the potential of LLMs to improve the construction of knowledge graphs ^11,12^. Trajanoska and colleagues introduced an NLP-based approach to build a Knowledge Graph on sustainability using web documents, showcasing that valuable information can be auto-extracted from unstructured data for decision-making and process modeling ^12^. Carta et. al. presented a novel method for knowledge graph construction using GPT-3.5, employing iterative zero-shot techniques, eliminating the need for external resources, and proving scalable and adaptable across diverse contexts ^11^. In terms of relation extraction in the clinical domains, Tang and colleagues assessed ChatGPT’s capability in clinical text mining by using ChatGPT-generated synthetic data. Their method achieved significant performance improvements over the state-of-the-art models and addressed data privacy concerns, highlighting a promising direction for LLMs in clinical contexts ^13^. Reichenpfader et. al. explored the use of large language models (LLMs) in extracting information from free-text radiology reports, a data source often overlooked for secondary uses like research ^**?**^. Torii and colleagues developed and compared three systems for extracting social determinants of health from clinical narratives using machine learning, LLM and hand-crafted rules ^14^. In a very recent work, Li et al. proposed a method called KnowCoder designed for Universal Information Extraction (UIE) through code generation, featuring a unique code-style schema representation to transform diverse schemas into Python classes for LLM-friendly processing ^15^.

Recently, there has been a shift towards more rigorous evaluation of LLMs in relation extraction tasks, with the introduction of new benchmarks focused on biological and biomedical datasets. Jahan et. al. presented the first extensive evaluation of LLMs in the biomedical domain, assessing 4 popular LLMs across 6 biomedical tasks and 26 datasets. Their findings revealed that zero-shot LLMs can surpass state-of-the-art models on datasets with smaller training sets, highlighting LLMs’ specialization potential in the biomedical field despite varying performance across tasks and their current limitations compared to models fine-tuned on larger datasets ^16^. Babaiha et. al. explored the use of OpenAI’s latest GPT models for one-shot extraction of Biological Expression Language (BEL) relations in biomedical knowledge graphs (KGs) and compared their efficacy against manually curated KGs by domain experts ^17^.

Our work distinguishes itself by focusing on the application of Large Language Models (LLMs) for specialized relation extraction tasks within the biomedical domain, specifically targeting the extraction of regulatory relations between genes, medications, and biological processes to augment biological knowledge graphs. Unlike existing studies that primarily explore general RE capabilities, few-shot learning, or synthetic data enhancement, our approach delves into leveraging LLMs for detailed and domain-specific knowledge extraction leveraging retrieval augmented generation, aiming to directly impact the enrichment of biomedical knowledge bases.

## 3 Methods

### 3.1 Corpus

To assess the capabilities of the LLMs in identifying such relationships, we use a manually annotated set of 104 titles extracted from PubMed articles. The titles were randomly selected from titles of all PubMed articles published in the last ten years that were tagged with the MeSH term “wound healing”. The gold standard corpus comprising of 104 titles feature at least one gene or medication concept and one GO biological concept. The biological process concepts are alligned with the GO vocabulary, while gene and medication annotations are linked automatically to the NCBI Gene database and RxNorm. Additionally, each title is annotated for regulatory relationships, represented by a (source, target, relation) triplet: the source being a gene or medication, the target being a biological process, and the relation classified as positive (upregulation) or negative (downregulation).

### 3.2 Experimental Framework

In Figure 1, we present an overview of the proposed biological process concept recognition, grounding, and relation extraction pipeline. In this subsection, we provide an in-depth description of each component within this pipeline, offering a comprehensive understanding of how each element contributes to the extraction of relationships between genes, medications, and biological process entities.

**Figure 1:**
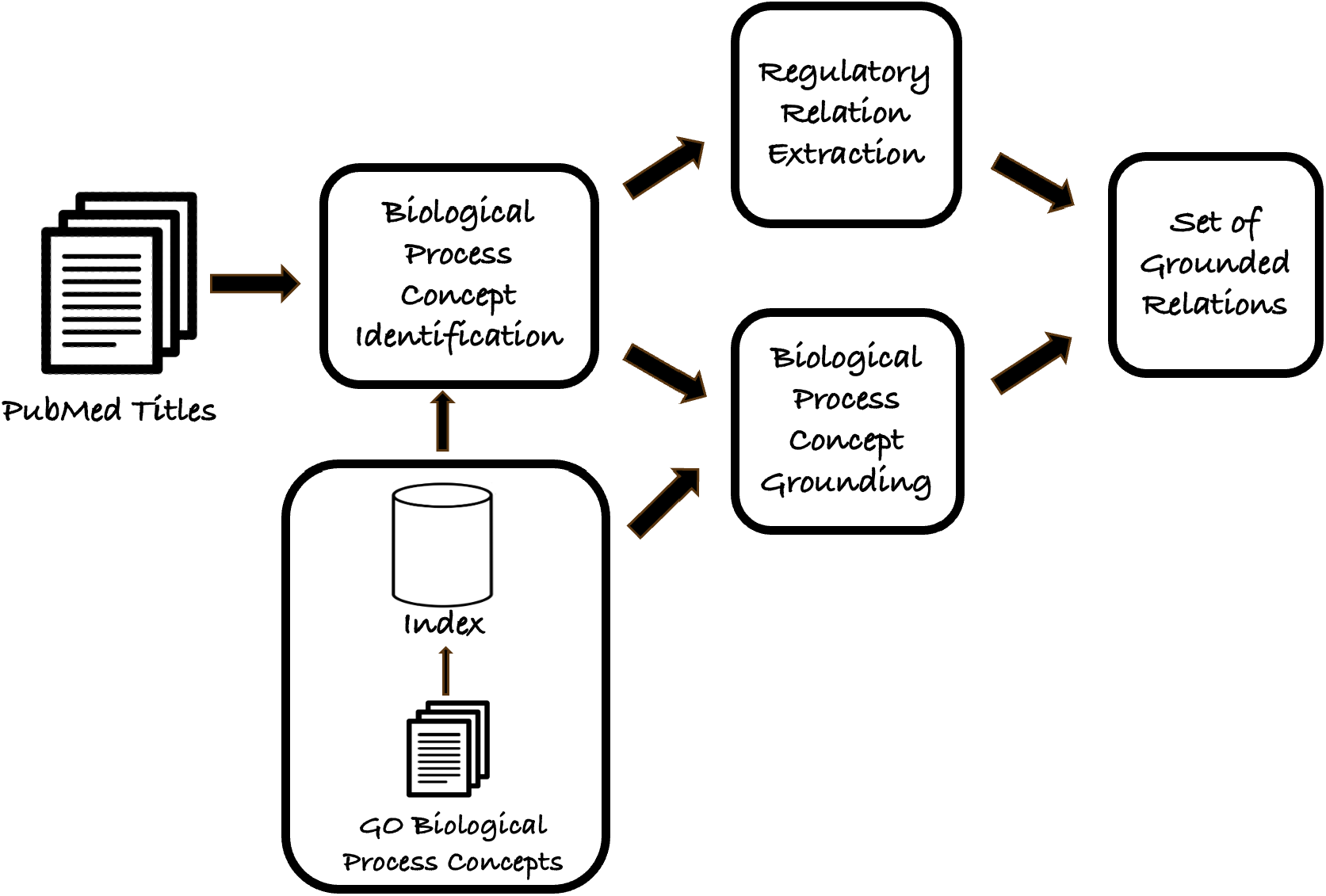
Biological process concept identification, concept grounding and relation extraction pipeline

#### 3.2.1 Biological Process Concept Identification

Over the years, there have been significant advancements in developing Named Entity Recognition (NER) tools that can efficiently identify gene and medication concepts within natural language texts. Tools such as “PubTator” and “scispaCy” have demonstrated proficiency in performing biomedical NER tasks and grounding identified biomedical concepts within existing knowledge bases. Since we utilize PubTator and scispaCy for the identification and grounding of gene and medication concepts, evaluating the NER capabilities of GPT-4 in identifying those concepts falls outside the scope of this study.

Our primary focus is on leveraging GPT-3.5 and GPT-4 in conjunction with the Retrieval Augmented Generation (RAG) technique to identify biological concepts in natural language texts. The RAG approach further enhances the LLMs’ advanced ability to generate more accurate, relevant, and contextually grounded annotations of biological processes. For the implementation of RAG, we constructed a vector database containing the names of GO biological process concepts. When a user query is made, the queried title is transformed into a vector representation and matched against this vector database to identify the most relevant *k* concepts based on the title. Subsequently, the RAG model enriches the input prompt of the LLMs by incorporating these pertinent GO concepts into the context. The implementation details of the RAG approach are described in Section 4.1. The NER prompt for GPT-3.5 and GPT-4 was developed following the methodologies outlined by Matentzoglu et al. and the prompting techniques of CurateGPT^‡ 18^. The query template used for the NER task is detailed in Appendix A.1.

#### 3.2.2 Biological Process Concept Grounding

Grounding natural language phrases to GO terms significantly enhances the understanding and categorization of biological processes, marking a substantial contribution to biomedical research. Traditional NER tools often face challenges in aligning biological processes expressed in natural language with standardized terms in knowledge bases. In our approach, we leverage both GPT-3.5 and GPT-4 with RAG-based prompting to effectively ground biological process concepts. For each entity phrase identified as a biological process during NER, grounding is performed individually by presenting the most relevant *k* concepts from the vector database, along with the title as additional context for more accurate grounding. Incorporating the title sentence into the grounding prompt aids the LLMs in more precise grounding of concepts. For instance, without any context, the entity phrase “cell proliferation” might be mapped to the more generic “cell population proliferation” concept within the GO vocabulary. However, when the title provides additional context, GPT-4 can try to assign the entity phrase to a more specific concept based on the context, such as “fibroblast proliferation” or “endothelial cell proliferation”. The prompt template for grounding biological concepts is elaborated in Appendix A.2.

#### 3.2.3 Regulatory Relation Extraction

In our study, we employ both GPT models for zero-shot relation extraction (RE) to identify (source, target, relation) triplets, where the source is represented by a gene or medication, the target by a biological process, and the relation indicates either positive (upregulation) or negative (downregulation). To mitigate the issues of hallucination and the extraction of an incomplete set of relations, we formulate the relation extraction task as a statement classification task. In this approach, we instruct the LLMs to classify a series of statements provided through user queries as *true, false*, or *unknown*, while the title sentence serves as the contextual backdrop for this classification task. Particularly, for each pair of gene or medication and biological process concepts, we generate two candidate statements indicating whether an upregulation or downregulation relationship exists between the two concepts. For instance, given the medication “Rapamycin” and the biological process “wound healing,” we would ask the LLMs to classify the statements “Rapamycin upregulates wound healing” and “Rapamycin downregulates wound healing” as *true, false*, or *unknown*, based on the associated title provided as context. Additionally, we request the LLMs to generate a chain-of-thought (CoT) reasoning for each statement in support of its classification. The statements identified as *true* are used to compile the final set of relations, from which we extract the (source, target, relation) triplets. The template for the RE prompt is provided in Appendix A.3.

## 4 Experimental Evaluation

We evaluate the performance of three key tasks: biological process concept identification, concept grounding, and relation extraction using the manually annotated corpus of 104 titles described in Section 3.1.

### 4.1 Evaluation Framework

In this study, we evaluate the performance of two most contemporary LLMs by OpenAI, namely GPT-3.5 and GPT-4, across the three tasks. For the tasks of biological process concept recognition and concept grounding, we employ **scispaCy** as a baseline model, drawing on its proven efficacy in biomedical NER and entity grounding tasks. Additionally, we implemented a rule-based relation extraction pipeline using **scispaCy** as a baseline for comparing relation extraction performance based on our previous work ^3^. This multifaceted approach allows us to assess the strengths and limitations of both traditional NLP tools and cutting-edge LLMs in handling the complexities of biological information extraction tasks.

#### Implementation Details

In the implementation of RAG for our study, we utilize LangChain^§^ to orchestrate an ensemble retriever strategy aimed at extracting the most relevant list of candidate Gene Ontology (GO) biological process concepts. This ensemble retriever combines the strengths of two distinct retrieval methods: a vector database retriever and a BM25 retriever. The vector database retriever operates by fetching the top 50 GO biological process concepts relevant to the queried title from a Chroma vector database. This database indexes the names of GO biological process concepts using embeddings generated by the “text-embedding-ada-002” embedding model of OpenAI. This method leverages the semantic understanding capabilities of the embeddings to find concepts that are contextually related to the query title. Conversely, the Okapi BM25 retriever employs a bag-of-words retrieval function, which ranks the GO concept names based on the occurrences of query terms within each concept name and returns the top 10 most relevant concepts. This approach is particularly effective for capturing direct term matches and is less influenced by the contextual nuances that vector-based methods excel at.

#### Evaluation Metrics

We evaluate all three tasks using traditional metrics such as precision, recall, and F1 score. For the NER task, it’s observed that both GPT models tend to annotate a broader span that encompasses the biological processes along with contextual words. For instance, the concept “wound healing” might be annotated as “cutaneous wound healing” or “dermal wound healing”. In this context, we deem an annotation correct if the ground truth concept annotation span is contained within the broader GPT annotation span. In the case of concept grounding, an exact match criterion is applied, comparing the model’s output directly against the ground truth annotations. This means that the grounded concept ID must precisely align with the established ground truth ID for it to be considered correct.

For the RE task, we define each relation as a triplet that includes a source (either a gene or a drug), a target (a biological process), and a relation (positive or negative, denoting the source’s effect on the target). The precision in this context is calculated as the fraction of correctly identified relations out of all the relations predicted by the model. Recall is measured as the fraction of correctly identified relations out of all actual relations present in the ground truth annotations. The F1 score is calculated as the harmonic mean of Precision and Recall, providing a single metric that balances both the precision and recall of the model’s predictions.

### 4.2 Results and Discussion

As we conduct the relation extraction (RE) task on a novel dataset that details the regulatory connections between genes or medications and biological functions, it is important to note that there are currently no existing models specifically tailored to address this particular task. Consequently, we face challenges in evaluating the performance of the LLMs against any established benchmarks or alternative models due to the absence of prior work in this area.

#### 4.2.1 Recognition and Grounding of Biological Processes Concepts

Table 1 presents the comparative performance of the baseline, GPT-3.5, and GPT-4 in the tasks of Named Entity Recognition (NER) of biological concepts and the subsequent grounding of these identified entities. Among the evaluated models, GPT-4 stands out by achieving the highest precision, recall, and F1 score in both the Named Entity Recognition (NER) and concept grounding tasks, surpassing the performance of GPT-3.5 and the baseline model. This underscores the effectiveness of GPT-4 in accurately identifying and grounding biological process concepts expressed in natural language, highlighting its advanced capabilities in handling the complexities of biomedical text analysis. GPT-3.5 also shows an improvement over the baseline in the NER task, achieving higher precision, recall, and F1 score. However, it occasionally misclassifies entities such as cell types (e.g., epithelial cells, endothelial cells) as biological process concepts. This misclassification is partly due to the fact that GO concept names sometimes include these cell names as substrings (e.g., endothelial cell proliferation, epithelial cell proliferation). The challenge is exacerbated when multiple biological process concepts are mentioned within a single sentence, which may also be attributed to the imperfect retrieval of candidate concepts through the RAG mechanism.

**Table 1:** Results for named entity recognition (NER) and entity grounding of biological process concepts in the manually annotated corpus introduced in Section 3.1.

| Model | Biological Process Concept Recognition |  |  | Biological Process Concept Grounding |  |  |
| --- | --- | --- | --- | --- | --- | --- |
|  | Precision | Recall | F1-Score | Precision | Recall | F1-Score |
| scispaCy | 0.71 | 0.59 | 0.64 | 0.57 | 0.46 | 0.51 |
| GPT-3.5 | 0.79 | 0.78 | 0.79 | 0.55 | 0.54 | 0.55 |
| GPT-4 | **0.89** | **0.93** | **0.91** | **0.71** | **0.74** | **0.72** |

In the concept grounding task, GPT-4 surpasses both GPT-3.5 and the baseline across all evaluation metrics. While the baseline model exhibits slightly higher precision than GPT-3.5, the latter achieves better recall and F1-score. However, GPT-4 stands out with much higher precision, recall, and F1-score, showcasing its ability to ground concepts to the most specific concept IDs in the GO knowledgebase by leveraging contextual information. One major drawback of the baseline model is its inability to utilize contextual information for concept grounding. For instance, in the title “Er:YAG laser promotes proliferation and wound healing capacity of human periodontal ligament fibroblasts through Galectin-7 induction,” GPT-4 correctly maps “proliferation” to “fibroblast proliferation” based on context, whereas the baseline model can only associate “proliferation” with a more generic GO concept like “cell proliferation.” It is also important to note that errors in the initial concept recognition phase are propagated to errors in concept grounding phase which comes next in the proposed pipeline (Figure 1). Such errors may also arise when a biological process term is closely associated with multiple concepts within the GO knowledgebase. For instance, the term “neovascularization” is closely related to both “vasculogenesis” and “angiogenesis” in the GO database. While the ground truth annotation maps “neovascularization” to “vasculogenesis”, we observe that both GPT-3.5 and GPT-4 map it to “angiogenesis”. This highlights the challenge of distinguishing between closely related concepts and the importance of precise initial recognition to ensure accurate grounding.

#### 4.2.2 Extraction and Ground of Regulatory Relations

Table 2 outlines the comparative performance of the baseline model, GPT-3.5, and GPT-4 in performing the relation extraction (RE) task. The results reveal that GPT-4 surpasses both the baseline and GPT-3.5 across all evaluation metrics. Despite GPT-3.5’s lower precision relative to the baseline, it compensates with significantly improved Recall and F1-score compared to the baseline. The baseline follows a rule-based approach for relation extraction and manages to identify a limited number of relations with high precision but suffers from notably low recall which is a common limitation of rule-based methods. This highlights the capability of Large Language Models (LLMs) to extract regulatory relations directly from natural language texts, bypassing the necessity for any specialized model training or fine-tuning. It is crucial to recognize that errors in relation extraction also propagate from prior stages, such as biological process concept recognition (Figure 1). GPT-4 showcases superior performance in the grounding of relations across all evaluation metrics. On the other hand, GPT-3.5 exhibits significant difficulties in this domain. This can be attributed to accumulated errors from preceding stages in the pipeline, such as concept recognition, grounding, and the initial extraction of relations. It’s crucial to understand that the models do not perform in relation grounding explicitly. Rather, the GO IDs derived from the process of biological concept grounding are utilized to establish the set of grounded relations.

**Table 2:** Results for extraction and grounding of regulatory relations between genes or medications and biological processes in the manually annotated corpus introduced in Section 3.1.

| Model | Relation Extraction |  |  | Relation Grounding |  |  |
| --- | --- | --- | --- | --- | --- | --- |
|  | Precision | Recall | F1-Score | Precision | Recall | F1-Score |
| scispaCy | 0.75 | 0.15 | 0.25 | 0.62 | 0.12 | 0.20 |
| GPT-3.5 | 0.63 | 0.73 | 0.68 | 0.44 | 0.51 | 0.47 |
| GPT-4 | **0.88** | **0.90** | **0.89** | **0.70** | **0.72** | **0.71** |

The comprehensive evaluation of the three tasks unequivocally demonstrates the superior performance of GPT-4 compared to GPT-3.5. This outcome is expected, considering GPT-4’s enhanced optimization, its training on more recent datasets, and its proven expertise in analyzing biological texts. The outputs generated by the model are available for review^¶^.

## 5 Limitations and Future Work

Even though our results demonstrate the superior performance of the LLMs, specifically GPT-4, across the three tasks of recognition and grounding of biological process concepts, and extraction of regulatory relations, there are some limitations to the proposed approach. The current approach utilizes the whole sentence for extracting a list of related GO biological process concepts for RAG based approach. This can be problematic for longer titles or texts because the similarity score is averaged which will result in imperfect extraction of candidate GO concepts. Chunking the texts with overlapping fixed length context window, and extracting candidate GO concepts based on the similarity across different context windows might be a better approach for retrieving more relevant candidate concepts. Another limitation lies in the area of relation extraction. The capabilities of LLMs in open-ended relation extraction is limited. In relation extraction, both GPT-3.5 and GPT-4 find it challenging to navigate through sentences with contextual ambiguities, complicating precise extraction of regulatory relations. This issue becomes prominent in instances where sentences contain many relations. To address this, we have designed RE prompts for the LLMs to assess user-generated candidate statements depicting such regulatory relations among concepts as true, false, or unknown. However, the complexity of this task increases significantly with the number of gene, medication, and biological process concepts involved. It’s important to note that none of the LLMs have been fine-tuned for the specific tasks outlined in this study, which can lead to somewhat inconsistent performance. This results in non-deterministic performance of the LLMs where the LLMs can generate different outputs even when the temperature parameter is set to 0.

In future endeavors, we plan to refine the RAG-based retrieval strategy by implementing text segmentation to more effectively identify candidate concepts for the annotation and grounding of biological processes. Additionally, we aim to improve prompt engineering to boost the performance of the LLMs across all three tasks: concept recognition, grounding, and relation extraction. Another key objective for future direction is to fine-tune LLMs specifically for the tasks outlined in this study in order to conduct a comparative analysis between the fine-tuned models and the approach proposed in our current work.

## 6 Conclusions

In this research, we developed a comprehensive pipeline utilizing Large Language Models (LLMs) for the purpose of relation extraction, aiming to uncover the impact of genes and medications on the biological processes involved in wound healing. The capabilities of both GPT-3.5 and GPT-4 were assessed through identification and grounding of biological processes and relation extraction tasks using a manually annotated corpus serving as the benchmark for evaluation. Our findings underscore the superior performance of GPT-4 over both its predecessor, GPT-3.5, and a baseline model across these tasks, highlighting the model’s advanced capabilities in handling the complexities of biomedical text analysis without the need for specific model training or fine-tuning. Despite these promising results, we also acknowledge the limitations of our current approach. The challenges identified pave the way for future improvements, including the refinement of the RAG-based retrieval approach and the exploration of more effective prompt engineering strategies. As we look forward to these developments, it is clear that the integration of LLMs into the workflow of biomedical research and drug discovery holds transformative potential. By enhancing the accuracy and efficiency of extracting and grounding biological concepts and regulatory relations, LLMs can significantly contribute to the acceleration of research advancements and the identification of novel therapeutic targets.

## Supporting information

Latex Source

### A Appendix

#### A.1 NER Prompt Template

Following is the list of candidate Gene Ontology (GO) biological process concepts:

{*candidates*}

Your job is to parse the following title and identify all instances of the same, equivalent, or similar biological process concepts given the above list of candidate concepts.

Mark up the concepts (if any) in double square brackets preserving the original text of the title inside the brackets like [[original text]].

If no biological process is explicitly mentioned in the supplied text, return the supplied text unchanged.

Example:

title: Interference with KCNJ2 inhibits proliferation, migration and EMT progression of papillary thyroid carcinoma cells by upregulating GNG2 expression.

output: Interference with KCNJ2 inhibits [[proliferation]], [[migration]] and EMT progression of papillary thyroid carcinoma cells by upregulating GNG2 expression.

title: {*text*}

output:

#### A.2 Grounding Prompt Template

Your role is to assign a concept ID that best matches the supplied text, using the supplied list of candidate concepts. Return as a string “CONCEPT NAME|CONCEPT ID|Score” triplet where the score represents the confidence of the assignment and should be between 0 and 1. Only use concept IDs from the supplied list of candidate concepts. Only return a row if the concept ID is a match for the input text. If there is no match, return NOT FOUND.

Here are the candidate concepts, as CONCEPT NAME|CONCEPT ID pairs: {*candidates*}

The overall context for this is the sentence: {*title*}

Concept to ground: {*text*}

#### A.3 RE Prompt Template

{*title*}

Based on the above statement only, your job is to label the following sentences as true, false, or unknown and provide a chain-of-thought (CoT) supporting the answer.

The answers should be in the following format.

1. label//CoT
2. label//CoT
3. label//Cot

…… etc.

{*candidate sentences*}

## Footnotes

† https://geneontology.org

‡ https://github.com/monarch-initiative/curate-gpt

§ https://www.langchain.com/

¶ https://github.com/juijayati/LLM-Based-NER-RE

## References

1. Crum RJ, Johnson SA, Jiang P, Jui JH, Zamora R, Cortes D, et al. Transcriptomic, proteomic, and morphologic characterization of healing in volumetric muscle loss. Tissue Engineering Part A. 2022;28(23-24):941–57.

2. Hosgood G. Stages of wound healing and their clinical relevance. Veterinary Clinics: Small Animal Practice. 2006;36(4):667–85.

3. Jui JH, Hauskrecht M. Uncovering the Effects of Genes, Proteins, and Medications on Functions of Wound Healing: A Dependency Rule-Based Text Mining Approach Leveraging GPT-4. In: 2023 IEEE EMBS International Conference on Biomedical and Health Informatics (BHI) (IEEE BHI 2023). Pittsburgh, USA; 2023. p. 3.

4. Fernández-Torras A, Duran-Frigola M, Bertoni M, Locatelli M, Aloy P. Integrating and formatting biomedical data as pre-calculated knowledge graph embeddings in the Bioteque. Nature Communications. 2022;13(1):5304.

5. Himmelstein DS, Lizee A, Hessler C, Brueggeman L, Chen SL, Hadley D, et al. Systematic integration of biomedical knowledge prioritizes drugs for repurposing. Elife. 2017;6:e26726.

6. Ioannidis VN, Song X, Manchanda S, Li M, Pan X, Zheng D, et al. DRKG - Drug Repurposing Knowledge Graph for Covid-19; 2020. https://github.com/gnn4dr/DRKG/.

7. Wadhwa S, Amir S, Wallace BC. Revisiting relation extraction in the era of large language models. In: Proceedings of the conference. Association for Computational Linguistics. Meeting. vol. 2023. NIH Public Access; 2023. p. 15566.

8. Xu X, Zhu Y, Wang X, Zhang N. How to Unleash the Power of Large Language Models for Few-shot Relation Extraction? In: The Fourth Workshop on Simple and Efficient Natural Language Processing; 2023. p. 190.

9. Xu B, Wang Q, Lyu Y, Dai D, Zhang Y, Mao Z. S2ynRE: Two-stage Self-training with Synthetic data for Low-resource Relation Extraction. In: Proceedings of the 61st Annual Meeting of the Association for Computational Linguistics (Volume 1: Long Papers); 2023. p. 8186–207.

10. Yuan C, Xie Q, Ananiadou S. Zero-shot Temporal Relation Extraction with ChatGPT. In: The 22nd Workshop on Biomedical Natural Language Processing and BioNLP Shared Tasksm; 2023. p. 92–102.

11. Carta S, Giuliani A, Piano L, Podda AS, Pompianu L, Tiddia SG. Iterative Zero-Shot LLM Prompting for Knowledge Graph Construction. arXiv preprint 230701128. 2023.

12. Trajanoska M, Stojanov R, Trajanov D. Enhancing Knowledge Graph Construction Using Large Language Models. arXiv preprint 230504676. 2023.

13. Tang R, Han X, Jiang X, Hu X. Does synthetic data generation of llms help clinical text mining? arXiv preprint 230304360. 2023.

14. Torii M, Finn IM, Doan S, Wang P, Yang EW, Zisook DS. Task formulation for extracting social determinants of health from clinical narratives. arXiv preprint 230111386. 2023.

15. Li Z, Zeng Y, Zuo Y, Ren W, Liu W, Su M, et al. KnowCoder: Coding Structured Knowledge into LLMs for Universal Information Extraction. arXiv preprint 240307969. 2024.

16. Jahan I, Laskar MTR, Peng C, Huang JX. A comprehensive evaluation of large language models on benchmark biomedical text processing tasks. Computers in Biology and Medicine. 2024:108189.

17. Babaiha NS, Rao SG, Klein J, Schultz B, Jacobs M, Hofmann-Apitius M. Rationalism in the face of GPT hypes: Benchmarking the output of large language models against human expert-curated biomedical knowledge graphs. Artificial Intelligence in the Life Sciences. 2024;5:100095.

18. Matentzoglu N, Caufield JH, Hegde HB, Reese JT, Moxon S, Kim H, et al. MapperGPT: Large Language Models for Linking and Mapping Entities. arXiv preprint 231003666. 2023.

