## Supplementary material for "Harnessing the Power of Large Language Models (LLMs) to Unravel the Influence of Genes and Medication on Biological Processes of Wound Healing^*^": Latex Source: amia.bib

%%
%% WinShell 3.3.1.10
%% http://www.winshell.org/
%%

% Strings


% Preamble


% BibTeX Entries
@article{gardner90,
 author               = {{Gardner RM, Golubjatnikov OK, Laub RM, Jacobson JT, Evans RS}},
 journal              = {Comput Biomed Res},
 pages                = {514-28},
 title                = {{Computer-critiqued blood ordering using the HELP system}},
 volume               = {23},
 year                 = {1990},
 }

@article{pryor83,
 author               = {{Pryor TA, Gardner RM, Clayton RD, Warner HR}},
 journal              = {J Med Sys},
 pages                = {87--101},
 title                = {{The HELP system}},
 volume               = {7},
 year                 = {1983},
 }
