## Supplementary material for "Harnessing the Power of Large Language Models (LLMs) to Unravel the Influence of Genes and Medication on Biological Processes of Wound Healing^*^": Latex Source: amia.pdf

### Title of American Medical Informatics Association Submission

**Firstname A. Lastname, MD, MPH[1], Firstname B. Lastname, MD, PhD[2]**
**[1] Institution, City, MA; [2]Institution, City, CA**

##### Abstract

*Abstract text goes here, justified and in italics. The abstract would normally be one paragraph long. See Table 1. for appropriate abstract length by submission type.*

##### Introduction

This template should be used as a starting point for AMIA submissions. A number of Word styles, all beginning with the word "AMIA", are available for use in your submissions.

It is important to review the AMIA Call for Participation where types of submissions considered and general requirements for each submission type are listed. All submissions must conform to the format and presentation requirements described in the CFP and at the submission site.

##### Another Major Heading and References

This sentence has two reference citations[1, 2].

More text of an additional paragraph, with a figure reference (Figure 1) and a figure inside a Word text box below. Figures need to be placed as close to the corresponding text as possible and not extend beyond one page.

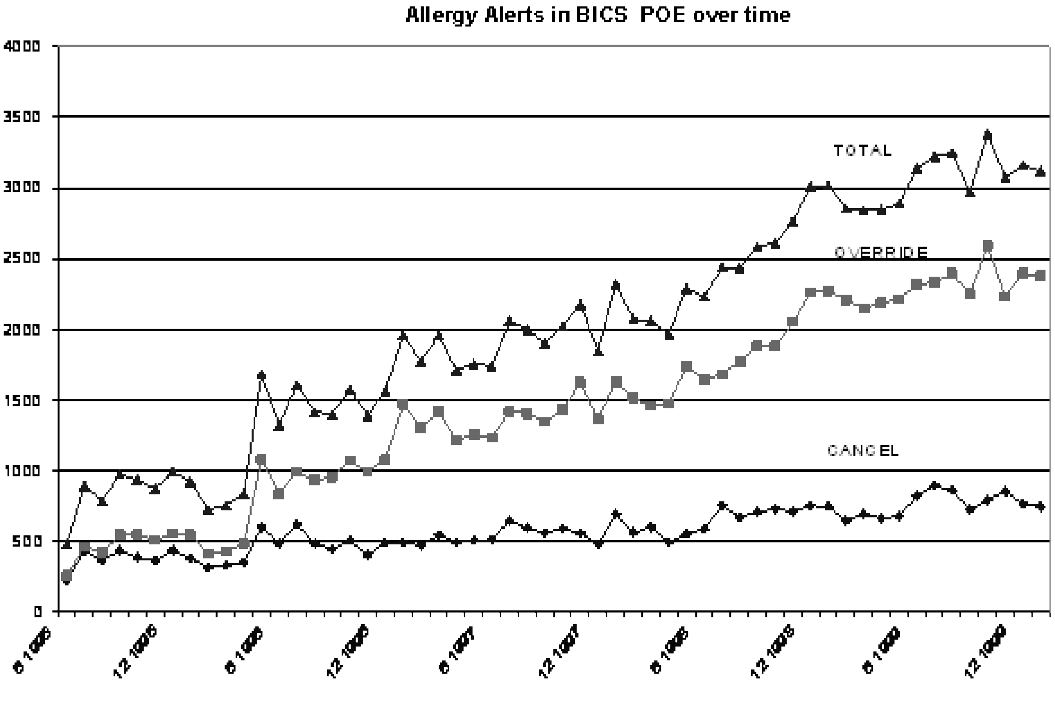


Figure 1: Total allergy alerts, overridden alerts, or drug order cancelled.

This is additional text added just to show the one-column formatting. This is additional text added just to show the one-column formatting. This is additional text added just to show the one-column formatting. This is additional text added just to show the one-column formatting. This is additional text added just to show the one-column formatting. This is additional text added just to show the one-column formatting. This is additional text added just to show the one-column formatting.

This paragraph contains a reference to a table just below (Table 1). All tables need to be placed as close to the corresponding text as possible, But each individual table should be on one page and not extend to multiple pages unless labeled as “Continued”.

| Submission Type | Abstract Length | Page Length Maximum |
|---|---|---|
| Paper | 125-150 words | Ten |
| Student Paper | 125-150 words | Ten |
| Poster | 50-75 words | One |
| Panel | 150-200 words | Three |
| Workshop | 150-200 words | Three |
| Theater-style Demonstrations | 150-200 words | One |
| Partnerships in Innovation | 150-200 words | Three |
| ACMI Senior Presentations | 150-200 words | Two |

Table 1: Submission type, abstract length, and page length maximum for AMIA submissions.

This is another paragraph.

**Sample paragraph heading** Lorem ipsum dolor sit amet, consectetuer adipiscing elit. Ut purus elit, vestibulum ut, placerat ac, adipiscing vitae, felis. Curabitur dictum gravida mauris. Nam arcu libero, nonummy eget, consectetuer id, vulputate a, magna. Donec vehicula augue eu neque. Pellentesque habitant morbi tristique senectus et netus et malesuada fames ac turpis egestas. Mauris ut leo. Cras viverra metus rhoncus sem. Nulla et lectus vestibulum urna fringilla ultrices. Phasellus eu tellus sit amet tortor gravida placerat. Integer sapien est, iaculis in, pretium quis, viverra ac, nunc. Praesent eget sem vel leo ultrices bibendum. Aenean faucibus. Morbi dolor nulla, malesuada eu, pulvinar at, mollis ac, nulla. Curabitur auctor semper nulla. Donec varius orci eget risus. Duis nibh mi, congue eu, accumsan eleifend, sagittis quis, diam. Duis eget orci sit amet orci dignissim rutrum.

#### Conclusion

Your conclusion goes at the end, followed by References. References begin below with a header that is centered. Only the first word of an article title is capitalized in the References.

**Acknowledgments** Lorem ipsum dolor sit amet, consectetuer adipiscing elit. Ut purus elit, vestibulum ut, placerat ac, adipiscing vitae, felis. Curabitur dictum gravida mauris. Nam arcu libero, nonummy eget, consectetuer id, vulputate a, magna. Donec vehicula augue eu neque. Pellentesque habitant morbi tristique senectus et netus et malesuada fames ac turpis egestas. Mauris ut leo. Cras viverra metus rhoncus sem. Nulla et lectus vestibulum urna fringilla ultrices. Phasellus eu tellus sit amet tortor gravida placerat. Integer sapien est, iaculis in, pretium quis, viverra ac, nunc. Praesent eget sem vel leo ultrices bibendum. Aenean faucibus. Morbi dolor nulla, malesuada eu, pulvinar at, mollis ac, nulla. Curabitur auctor semper nulla. Donec varius orci eget risus. Duis nibh mi, congue eu, accumsan eleifend, sagittis quis, diam. Duis eget orci sit amet orci dignissim rutrum.

#### References

1. Pryor TA, Gardner RM, Clayton RD, Warner HR. The HELP system. J Med Sys. 1983;7:87–101.
2. Gardner RM, Golubjatnikov OK, Laub RM, Jacobson JT, Evans RS. Computer-critiqued blood ordering using the HELP system. Comput Biomed Res. 1990;23:514–28.
