## Supplementary material for "Harnessing the Power of Large Language Models (LLMs) to Unravel the Influence of Genes and Medication on Biological Processes of Wound Healing^*^": Latex Source: amia.tex

\documentclass{amia}
\usepackage{lipsum} %Remove if not needed
\setlength{\bibsep}{0pt} %Comment out if you don't want to condense the bibliography

\begin{document}


\title{Title of American Medical Informatics Association Submission}

\author{Firstname A. Lastname, MD, MPH$^1$, Firstname B. Lastname, MD, PhD$^2$ }

\institutes{
    $^1$ Institution, City, MA; $^2$Institution, City, CA
}

\maketitle

\section*{Abstract}

\textit{Abstract text goes here, justified and in italics.  The abstract would normally be one paragraph long.  See Table~\ref{tab:submission}. for appropriate abstract length by submission type.}

\section*{Introduction}

This template should be used as a starting point for AMIA submissions.  A number of Word styles, all beginning with the word ``AMIA", are available for use in your submissions.

It is important to review the AMIA Call for Participation where types of submissions considered and general requirements for each submission type are listed. All submissions must conform to the format and presentation requirements described in the CFP and at the submission site.


\section*{Another Major Heading and References}

This sentence has two reference citations\cite{pryor83,gardner90}.

More text of an additional paragraph, with a figure reference (Figure~\ref{fig:amia_figure}) and a figure inside a Word text box below.  Figures need to be placed as close to the corresponding text as possible and not extend beyond one page.

\begin{figure}[H]
  \centering
    \includegraphics[width=1.0\textwidth]{amia_figure1}
 \caption{Total allergy alerts, overridden alerts, or drug order cancelled.}
 \label{fig:amia_figure}
\end{figure}

This is additional text added just to show the one-column formatting. This is additional text added just to show the one-column formatting. This is additional text added just to show the one-column formatting.  This is additional text added just to show the one-column formatting. This is additional text added just to show the one-column formatting. This is additional text added just to show the one-column formatting.  This is additional text added just to show the one-column formatting.

This paragraph contains a reference to a table just below (Table~\ref{tab:submission}).  All tables need to be placed as close to the corresponding text as possible, But each individual table should be on one page and not extend to multiple pages unless labeled as ``Continued".

\begin{table}[H]
\begin{center}
\begin{tabular}{|l|l|l|}
\hline
Submission Type	            &Abstract Length	&Page Length Maximum \\
\hline
Paper                       &125-150 words	    &Ten                 \\
\hline 
Student Paper	            &125-150 words	    &Ten                 \\
\hline
Poster                      &50-75 words        &One                 \\
\hline
Panel	                    &150-200 words      &Three               \\
\hline
Workshop	                &150-200 words      &Three               \\
\hline
Theater-style Demonstrations&150-200 words	    &One                 \\
\hline
Partnerships in Innovation	&150-200 words	    &Three               \\
\hline
ACMI Senior Presentations	&150-200 words	    &Two                 \\
\hline
\end{tabular}
\end{center}
\caption{Submission type, abstract length, and page length maximum for AMIA submissions.}
\label{tab:submission}
\end{table}

This is another paragraph.

\paragraph{Sample paragraph heading}\lipsum[1]

\subsection*{Conclusion}

Your conclusion goes at the end, followed by References.  References begin below with a header that is centered.  Only the first word of an article title is capitalized in the References.

\subparagraph{Acknowledgments}\lipsum[1]

% References as numbers
\makeatletter
\renewcommand{\@biblabel}[1]{\hfill #1.}
\makeatother

% unstr is used to keep citation order
\bibliographystyle{vancouver}
\bibliography{amia}  

\end{document}
