## Supplementary material for "Harnessing the Power of Large Language Models (LLMs) to Unravel the Influence of Genes and Medication on Biological Processes of Wound Healing^*^": Latex Source: main.tex

\documentclass{amia}
\usepackage{lipsum} %Remove if not needed
\setlength{\bibsep}{0pt} %Comment out if you don't want to condense the bibliography
%\usepackage{graphicx}
\usepackage{amsmath}
\usepackage[hidelinks]{hyperref}
\usepackage{multirow}
\usepackage{verbatim}
\usepackage[symbol]{footmisc}
\usepackage{enumitem}
\usepackage{comment}
\usepackage{tcolorbox}
\usepackage{tabularray}
\hypersetup{colorlinks=true, allcolors=blue}

%\urlstyle{same}

\begin{document}

\pagenumbering{arabic}

\title{Harnessing the Power of Large Language Models (LLMs) to Unravel the Influence of Genes and Medication on Biological Processes of Wound Healing\footnote{Supported by the Defense Advanced Research Projects Agency (DARPA) through Cooperative Agreement D20AC00002 awarded by the U.S. Department of the Interior, Interior Business Center. The content of the article does not necessarily reflect the position or the policy of the Government, and no official endorsement should be inferred.}}

\author{Jayati H. Jui, MS, Milos Hauskrecht, PhD}

\institutes{
    University of Pittsburgh, PA 15260, USA\\
    \{jaj146,milos\}@pitt.edu
}

\maketitle

\section*{Abstract}

\textit{Recent advancements in Large Language Models (LLMs) have ushered in a new era for knowledge extraction in the
domains of biological and clinical natural language processing (NLP). In this research, we present a novel
approach to understanding the regulatory effects of genes and medications on biological processes central to wound
healing. Utilizing the capabilities of Generative Pre-trained Transformer (GPT) models by OpenAI, specifically GPT-3.5 and GPT-4, we developed a comprehensive pipeline for the identification and grounding of biological processes and the extraction of such regulatory relations. The performances of both GPTs were rigorously evaluated against a manually annotated corpus of 104 PubMed titles, focusing on their ability to accurately identify and ground biological process concepts and extract relevant regulatory relationships from the text. Our findings demonstrate that GPT-4, in particular, exhibits superior performance in all the tasks, showcasing its potential to facilitate significant advancements in biomedical research without requiring model fine-tuning. 
}
\input{sections/introduction}
\input{sections/background}
\input{sections/methods}
\input{sections/results}
\input{sections/limitations}
\input{sections/conclusions}

% References as numbers
\makeatletter
\renewcommand{\@biblabel}[1]{\hfill #1.}
\makeatother

% unstr is used to keep citation order
\bibliographystyle{vancouver}
\bibliography{mybibliography}  
\appendix
\input{sections/Appendix}
\end{document}
