## Supplementary material for "Harnessing the Power of Large Language Models (LLMs) to Unravel the Influence of Genes and Medication on Biological Processes of Wound Healing^*^": Latex Source: mybibliography.bib

@inproceedings{wadhwa2023revisiting,
  title={Revisiting relation extraction in the era of large language models},
  author={Wadhwa, Somin and Amir, Silvio and Wallace, Byron C},
  booktitle={Proceedings of the conference. Association for Computational Linguistics. Meeting},
  volume={2023},
  pages={15566},
  year={2023},
  organization={NIH Public Access}
}

@inproceedings{xu2023s2ynre,
  title={S2ynRE: Two-stage Self-training with Synthetic data for Low-resource Relation Extraction},
  author={Xu, Benfeng and Wang, Quan and Lyu, Yajuan and Dai, Dai and Zhang, Yongdong and Mao, Zhendong},
  booktitle={Proceedings of the 61st Annual Meeting of the Association for Computational Linguistics (Volume 1: Long Papers)},
  pages={8186--8207},
  year={2023}
}

@article{rajpoot2023gpt,
  title={GPT-FinRE: In-context Learning for Financial Relation Extraction using Large Language Models},
  author={Rajpoot, Pawan Kumar and Parikh, Ankur},
  journal={arXiv preprint arXiv:2306.17519},
  year={2023}
}

@inproceedings{yuan2023zero,
  title={Zero-shot Temporal Relation Extraction with ChatGPT},
  author={Yuan, Chenhan and Xie, Qianqian and Ananiadou, Sophia},
  booktitle={The 22nd Workshop on Biomedical Natural Language Processing and BioNLP Shared Tasks},
  pages={92--102},
  year={2023}
}

@article{carta2023iterative,
  title={Iterative Zero-Shot LLM Prompting for Knowledge Graph Construction},
  author={Carta, Salvatore and Giuliani, Alessandro and Piano, Leonardo and Podda, Alessandro Sebastian and Pompianu, Livio and Tiddia, Sandro Gabriele},
  journal={arXiv preprint arXiv:2307.01128},
  year={2023}
}

@article{trajanoska2023enhancing,
  title={Enhancing Knowledge Graph Construction Using Large Language Models},
  author={Trajanoska, Milena and Stojanov, Riste and Trajanov, Dimitar},
  journal={arXiv preprint arXiv:2305.04676},
  year={2023}
}

@inproceedings{xu2023unleash,
  title={How to Unleash the Power of Large Language Models for Few-shot Relation Extraction?},
  author={Xu, Xin and Zhu, Yuqi and Wang, Xiaohan and Zhang, Ningyu},
  booktitle={The Fourth Workshop on Simple and Efficient Natural Language Processing},
  pages={190},
  year={2023}
}


@article{tang2023does,
  title={Does synthetic data generation of llms help clinical text mining?},
  author={Tang, Ruixiang and Han, Xiaotian and Jiang, Xiaoqian and Hu, Xia},
  journal={arXiv preprint arXiv:2303.04360},
  year={2023}
}

@article{reichenpfader2023protocol,
  title={Protocol: Large language model-based information extraction from free-text radiology reports: a scoping review protocol},
  author={Reichenpfader, Daniel and M{\"u}ller, Henning and Denecke, Kerstin},
  journal={BMJ open},
  volume={13},
  number={12},
  year={2023},
  publisher={BMJ Publishing Group}
}

@article{torii2023task,
  title={Task formulation for extracting social determinants of health from clinical narratives},
  author={Torii, Manabu and Finn, Ian M and Doan, Son and Wang, Paul and Yang, Elly W and Zisook, Daniel S},
  journal={arXiv preprint arXiv:2301.11386},
  year={2023}
}

@article{crum2022transcriptomic,
  title={Transcriptomic, proteomic, and morphologic characterization of healing in volumetric muscle loss},
  author={Crum, Raphael J and Johnson, Scott A and Jiang, Peng and Jui, Jayati H and Zamora, Ruben and Cortes, Devin and Kulkarni, Mangesh and Prabahar, Archana and Bolin, Jennifer and Gann, Eric and others},
  journal={Tissue Engineering Part A},
  volume={28},
  number={23-24},
  pages={941--957},
  year={2022},
  publisher={Mary Ann Liebert, Inc., publishers 140 Huguenot Street, 3rd Floor New~…}
}

@article{hosgood2006stages,
  title={Stages of wound healing and their clinical relevance},
  author={Hosgood, Giselle},
  journal={Veterinary Clinics: Small Animal Practice},
  volume={36},
  number={4},
  pages={667--685},
  year={2006},
  publisher={Elsevier}
}

@article{quan2016multichannel,
  title={Multichannel convolutional neural network for biological relation extraction},
  author={Quan, Chanqin and Hua, Lei and Sun, Xiao and Bai, Wenjun and others},
  journal={BioMed research international},
  volume={2016},
  year={2016},
  publisher={Hindawi}
}

@article{verga2018simultaneously,
  title={Simultaneously self-attending to all mentions for full-abstract biological relation extraction},
  author={Verga, Patrick and Strubell, Emma and McCallum, Andrew},
  journal={arXiv preprint arXiv:1802.10569},
  year={2018}
}

@article{kim2018relation,
  title={Relation extraction for biological pathway construction using node2vec},
  author={Kim, Munui and Baek, Seung Han and Song, Min},
  journal={BMC bioinformatics},
  volume={19},
  pages={75--84},
  year={2018},
  publisher={Springer}
}

@article{ravikumar2017belminer,
  title={BELMiner: adapting a rule-based relation extraction system to extract biological expression language statements from bio-medical literature evidence sentences},
  author={Ravikumar, KE and Rastegar-Mojarad, Majid and Liu, Hongfang},
  journal={Database},
  volume={2017},
  pages={baw156},
  year={2017},
  publisher={Oxford University Press}
}

@article{wang2021nero,
  title={NERO: a biomedical named-entity (recognition) ontology with a large, annotated corpus reveals meaningful associations through text embedding},
  author={Wang, Kanix and Stevens, Robert and Alachram, Halima and Li, Yu and Soldatova, Larisa and King, Ross and Ananiadou, Sophia and Schoene, Annika M and Li, Maolin and Christopoulou, Fenia and others},
  journal={NPJ systems biology and applications},
  volume={7},
  number={1},
  pages={38},
  year={2021},
  publisher={Nature Publishing Group UK London}
}

@article{neumann2019scispacy,
  title={ScispaCy: fast and robust models for biomedical natural language processing},
  author={Neumann, Mark and King, Daniel and Beltagy, Iz and Ammar, Waleed},
  journal={arXiv preprint arXiv:1902.07669},
  year={2019}
}

@inproceedings{Jui2310:Uncovering,
AUTHOR    = {Jayati Halder Jui and Milos Hauskrecht},
TITLE    = {Uncovering the Effects of Genes, Proteins, and Medications on Functions of
Wound Healing: A Dependency {Rule-Based} Text Mining Approach Leveraging
{GPT-4}},
BOOKTITLE    = {2023 IEEE EMBS International Conference on Biomedical and Health
Informatics (BHI) (IEEE BHI 2023)},
ADDRESS    = {Pittsburgh, USA},
PAGES    = {3},
DAYS    = {15},
MONTH    = {October},
YEAR    = {2023}
}

@article{hetionet,
  title={Systematic integration of biomedical knowledge prioritizes drugs for repurposing},
  author={Himmelstein, Daniel Scott and Lizee, Antoine and Hessler, Christine and Brueggeman, Leo and Chen, Sabrina L and Hadley, Dexter and Green, Ari and Khankhanian, Pouya and Baranzini, Sergio E},
  journal={Elife},
  volume={6},
  pages={e26726},
  year={2017},
  publisher={eLife Sciences Publications, Ltd}
}

@misc{drkg2020,
  author = {Ioannidis, Vassilis N. and Song, Xiang and Manchanda, Saurav and Li, Mufei and Pan, Xiaoqin
            and Zheng, Da and Ning, Xia and Zeng, Xiangxiang and Karypis, George},
  title = {DRKG - Drug Repurposing Knowledge Graph for Covid-19},
  howpublished = "\url{https://github.com/gnn4dr/DRKG/}",
  year = {2020}
}

@misc{bc4eva,
  title={Proceedings of the fourth BioCreative challenge evaluation workshop},
  author={Arighi, CN and Cohen, KB and Hirschman, L and Lu, Z and Tudor, CO and Wiegers, T and Wilbur, WJ and Wu, CH},
  year={2013},
  publisher={Bethesda MD USA}
}

@article{bioteque,
  title={Integrating and formatting biomedical data as pre-calculated knowledge graph embeddings in the Bioteque},
  author={Fern{\'a}ndez-Torras, Adri{\`a} and Duran-Frigola, Miquel and Bertoni, Martino and Locatelli, Martina and Aloy, Patrick},
  journal={Nature Communications},
  volume={13},
  number={1},
  pages={5304},
  year={2022},
  publisher={Nature Publishing Group UK London}
}

@article{mappergpt,
  title={MapperGPT: Large Language Models for Linking and Mapping Entities},
  author={Matentzoglu, Nicolas and Caufield, J Harry and Hegde, Harshad B and Reese, Justin T and Moxon, Sierra and Kim, Hyeongsik and Harris, Nomi L and Haendel, Melissa A and Mungall, Christopher J},
  journal={arXiv preprint arXiv:2310.03666},
  year={2023}
}

@article{li2024knowcoder,
  title={KnowCoder: Coding Structured Knowledge into LLMs for Universal Information Extraction},
  author={Li, Zixuan and Zeng, Yutao and Zuo, Yuxin and Ren, Weicheng and Liu, Wenxuan and Su, Miao and Guo, Yucan and Liu, Yantao and Li, Xiang and Hu, Zhilei and others},
  journal={arXiv preprint arXiv:2403.07969},
  year={2024}
}

@article{jahan2024comprehensive,
  title={A comprehensive evaluation of large language models on benchmark biomedical text processing tasks},
  author={Jahan, Israt and Laskar, Md Tahmid Rahman and Peng, Chun and Huang, Jimmy Xiangji},
  journal={Computers in Biology and Medicine},
  pages={108189},
  year={2024},
  publisher={Elsevier}
}

@article{babaiha2024rationalism,
  title={Rationalism in the face of GPT hypes: Benchmarking the output of large language models against human expert-curated biomedical knowledge graphs},
  author={Babaiha, Negin Sadat and Rao, Sathvik Guru and Klein, J{\"u}rgen and Schultz, Bruce and Jacobs, Marc and Hofmann-Apitius, Martin},
  journal={Artificial Intelligence in the Life Sciences},
  volume={5},
  pages={100095},
  year={2024},
  publisher={Elsevier}
}

@article{caufieldstructured,
  title={Structured Prompt Interrogation and Recursive Extraction of Semantics (SPIRES): a method for populating knowledge bases using zero-shot learning},
  author={Caufield, J Harry and Hegde, Harshad and Emonet, Vincent and Harris, Nomi L and Joachimiak, Marcin P and Matentzoglu, Nicolas and Kim, HyeongSik and Moxon, Sierra and Reese, Justin T and Haendel, Melissa A and others},
  journal={Bioinformatics (Oxford, England)},
  pages={btae104}
}
