## Supplementary material for "Harnessing the Power of Large Language Models (LLMs) to Unravel the Influence of Genes and Medication on Biological Processes of Wound Healing^*^": Latex Source: readme.txt

AMIA Latex Template (updated 2016)
------------------

************
This is the latex template for American Medical Informatics Association by Antonio Jimeno Yepes. 
It has been updated from the original version authored by him. 
Modifications has been made to original cls file to address the compilation of the bibliography and some minor additions for the style.
- Tuan Amith
************

Essential Files:

* amia.cls is a latex class file to produce a PDF in the format required by AMIA.

* amia.tex is a latex document which can be used as example of use of amia.cls.
  In addition, the text in this file contains details about the AMIA format requirements.

  If you use dvipdf to produce the final PDF, margins might not follow the instructions of the template.
  You are advised to use "dvipdfm -p letter" instead.

* amia.bib is the latex bibliography file
  It is important to keep the bibliography style "unsrt" so the references follow the citation order.
