## Supplementary material for "Harnessing the Power of Large Language Models (LLMs) to Unravel the Influence of Genes and Medication on Biological Processes of Wound Healing^*^": Latex Source: Appendix.tex

\section{Appendix}
\subsection{NER Prompt Template}\label{ner_prompt}
\begin{tcolorbox}
Following is the list of candidate Gene Ontology (GO) biological process concepts:\\
$\{candidates\}$\\
Your job is to parse the following title and identify all instances of the same, equivalent, or similar biological process concepts given the above list of candidate concepts.\\
Mark up the concepts (if any) in double square brackets preserving the original text of the title inside the brackets like [[original text]].\\
If no biological process is explicitly mentioned in the supplied text, return the supplied text unchanged.\\ \\
Example:\\
title: Interference with KCNJ2 inhibits proliferation, migration and EMT progression of papillary thyroid carcinoma cells by upregulating GNG2 expression.\\
output: Interference with KCNJ2 inhibits [[proliferation]], [[migration]] and EMT progression of papillary thyroid carcinoma cells by upregulating GNG2 expression.
\\ \\
title: $\{text\}$\\
output:
\end{tcolorbox}

\subsection{Grounding Prompt Template}\label{gr_prompt}
\begin{tcolorbox}
Your role is to assign a concept ID that best matches the supplied text, using the supplied list of candidate concepts. Return as a string ``CONCEPT\_NAME$\vert$CONCEPT\_ID$\vert$Score'' triplet where the score represents the confidence 
of the assignment and should be between 0 and 1. Only use concept IDs from the supplied list of candidate concepts. Only return a row if the concept ID is a match for the input text. If there is no match, return NOT FOUND.\\ \\
Here are the candidate concepts, as CONCEPT\_NAME$\vert$CONCEPT\_ID pairs: $\{candidates\}$\\ \\
The overall context for this is the sentence: $\{title\}$\\ \\
Concept to ground: $\{text\}$
\end{tcolorbox}

\subsection{RE Prompt Template}\label{re_prompt}
%\subsubsection{Template}
\begin{tcolorbox}
$\{title\}$\\
Based on the above statement only, your job is to label the following sentences as true, false, or unknown
and provide a chain-of-thought (CoT) supporting the answer. \\
The answers should be in the following format.\\
1. label//CoT\\
2. label//CoT\\
3. label//Cot\\
..... etc.\\
$\{candidate \ sentences\}$
\end{tcolorbox}
