## Supplementary material for "Harnessing the Power of Large Language Models (LLMs) to Unravel the Influence of Genes and Medication on Biological Processes of Wound Healing^*^": Latex Source: tab_ner.tex

% \usepackage{tabularray}
\begin{table}
\centering
\begin{tblr}{
  cell{1}{1} = {r=2}{},
  cell{1}{2} = {c=5}{},
  cell{1}{7} = {c=5}{},
  cell{1-5}{1-11} = {c},
  vline{2-3} = {1}{},
  vline{7} = {2}{},
  vline{2,7} = {1-5}{},
  hline{1,3-6} = {-}{},
  hline{2} = {2-11}{},
}
Model    & Biological Process Concept Recognition &  &        &  &          & Biological Process Concept Grounding &  &        &  &          \\
         & Precision               &  & Recall &  & F1-Score & Precision            &  & Recall &  & F1-Score \\
scispaCy & 0.71                    &  & 0.59   &  & 0.64     & 0.57                 &  & 0.46   &  & 0.51     \\
GPT-3.5  & 0.79                     &  & 0.78    &  & 0.79      & 0.55                  &  & 0.54    &  & 0.55      \\
GPT-4    & \textbf{0.89}                     &  & \textbf{0.93}    &  & \textbf{0.91}      & \textbf{0.71}                  &  & \textbf{0.74}    &  & \textbf{0.72}      
\end{tblr}
\caption{Results for named entity recognition (NER) and entity grounding of biological process concepts in the manually annotated corpus introduced in Section \ref{corpus}}
\label{tab:ner}
\end{table}
