## Supplementary material for "Harnessing the Power of Large Language Models (LLMs) to Unravel the Influence of Genes and Medication on Biological Processes of Wound Healing^*^": Latex Source: tab_ner_new.tex

% \usepackage{tabularray}
\begin{table}
\centering
\caption{NER }
\label{tab:ner}
\begin{tblr}{
  cell{1}{1} = {r=2}{c},
  cell{1}{2} = {c=5}{c},
  vline{2-3} = {1}{},
  vline{7} = {2}{},
  vline{2,7} = {1-5}{},
  hline{1,3-6} = {-}{},
  hline{2} = {2-6}{},
}
Model    & NER (without grounding) &  &        &  &          \\
         & Precision               &  & Recall &  & F1-Score \\
scispaCy & 0.65                    &  & 0.65   &  & 0.65     \\
GPT-3.5  & 0.7                     &  & 0.7    &  & 0.7      \\
GPT-4    & 0.8                     &  & 0.7    &  & 0.8      
\end{tblr}
\end{table}
