## Supplementary material for "Harnessing the Power of Large Language Models (LLMs) to Unravel the Influence of Genes and Medication on Biological Processes of Wound Healing^*^": Latex Source: tab_re.tex

% \usepackage{tabularray}
\begin{table}
\centering
\begin{tblr}{
  cell{1}{1} = {r=2}{},
  cell{1}{2} = {c=5}{},
  cell{1}{7} = {c=5}{},
  cell{1-5}{1-11} = {c},
  vline{2-3} = {1}{},
  vline{7} = {2}{},
  vline{2,7} = {1-5}{},
  hline{1,3-6} = {-}{},
  hline{2} = {2-11}{},
}
Model    & Relation Extraction &  &        &  &          & Relation Grounding &  &        &  &          \\
         & Precision               &  & Recall &  & F1-Score & Precision            &  & Recall &  & F1-Score \\
scispaCy & 0.75                    &  & 0.15   &  & 0.25     & 0.62                 &  & 0.12   &  & 0.20     \\
GPT-3.5  & 0.63                     &  & 0.73    &  & 0.68      & 0.44                  &  & 0.51    &  & 0.47      \\
GPT-4    & \textbf{0.88}                     &  & \textbf{0.90}    &  & \textbf{0.89}      & \textbf{0.70}                  &  & \textbf{0.72}    &  & \textbf{0.71}      
\end{tblr}
\caption{Results for extraction and grounding of regulatory relations between genes or medications and biological processes in the manually annotated corpus introduced in Section \ref{corpus}}
\label{tab:re}
\end{table}
