## Supplementary figures and images for "Harnessing the Power of Large Language Models (LLMs) to Unravel the Influence of Genes and Medication on Biological Processes of Wound Healing^*^"

### flowchart.png

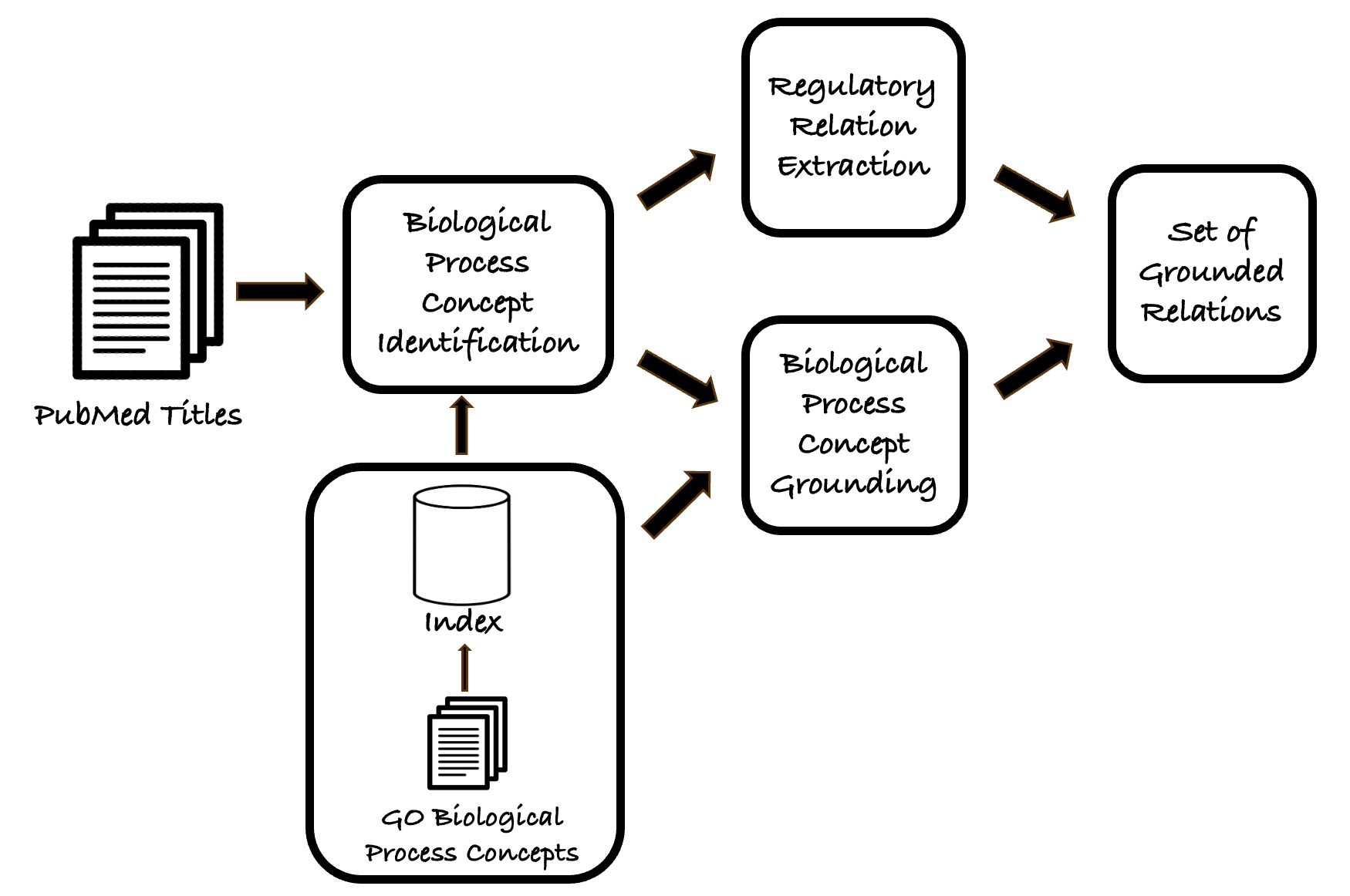
